# Automatic crystal identification for crystallography: *a comparison between direct methods and artificial intelligence strategies*

**DOI:** 10.1101/2024.08.22.609235

**Authors:** Camila F A Lages, Vitoria Y Nicoleti, Maicon R Correa, Paulo Mausbach, Felipe C Ramos, Evandro A de Araujo, Andrey Z Nascimento, Eduardo X Miqueles

## Abstract

Crystallography is an important, and well-established technique for assessing crystalline atomic structure of a wide variety of materials. An auxiliary optical microscope instrument is used to perform the data acquisition at the manaca beamline, a crystallography experimental station from sirius (the 4th generation Brazilian synchrotron). The importance of detecting these crystals in the auxiliary image is to define the best order in which each crystal will be measured by the incident beam since each one of them must be measured only once; improving beamtime and maximizing user experience at the facility. Detecting rough crystal positions from these auxiliary images is a scientific computing task, which could benefit from some open artificial intelligence tools. This manuscript compares a real-time object detection interface (Yolo) with a new, simple and effective strategy obtained from a particular partial differential equation.

## 1 Introduction

Crystallography is the experimental science of determining the arrangement of atoms in samples. It is fundamental in materials science, solid-state physics and structural biology [1]. At sirius (the 4th generation Brazilian synchrotron), during beamtime the user typically needs to manually select the crystals in a countable set (usually a sample consisting of a drop and protein crystals immersed in it). This selection is done using the help of several computational tools [2].

These crystals have different shapes and sizes and can be distributed randomly within the sample holder. The samples consist of crystals immersed in a drop of some liquid, depending on the condition of this drop, the crystal edges may be more or less defined in the obtained microscopic image. Another issue is the overlapping of crystals, a particular case that must be considered when identifying their centers. Usually, the beamline scientist will discard centers in crystal clusters since they can interfere when measuring the diffraction pattern of just one crystal. We also need to deal with background texture, noises, or salt crystals.

It is up to the beamline operating user to choose the crystal in the image (a drag-clicking operation) and inform its central position in pixel units. These pixel units are converted to motor positions, and the sample is then positioned to coincide with the incident beam. Although tempting, we do not need to find every crystal in the image, and a percentage of 80% is far enough for beamline users. The number of crystals within the sample drop is also unknown, being completely random.

In this work, we are interested in determining the largest number of crystals in the auxiliary microscopic image, comparing a specific artificial intelligence tool with a direct method. The time to carry out the crystals’ center identification should be low, as this is considered a pre-measurement process. The manual process of identifying crystals takes roughly 30 minutes, depending on the number of crystals available in the sample drop. We pose the challenge as the following Problem 1.

### Problem 1

*Find an image processing algorithm to identify and extract the maximum number of crystals from a microscope image. Let M be a n × n pixels discretized microscope image, we want to define a mapping C*: *providing m (number of detected crystals) pair of pixels p. We can define this as a mathematical operation p* = *C*[*M*].

The problem 1 was posed during an Integrative Think Tank initiative [3] in 2023 for an audience of graduate students and researchers of different scientific computing areas. The proposed partial differential equation, which provides us with a *direct* method was compared with the open tool called yolo (You only look once) [4]. Since this is not a manuscript with extensive mathematical descriptions, we avoid dense jargon - providing a more intuitive description of the problems and the obtained solutions. Both strategies are now integrated with the scanning routines at the manaca beamline [5] and the source code was made available for users.

## 2 Material and methods

Crystal identification on raw microscope images follows two main research branches: detection and segmentation. For the segmentation branch, two paths are possible, the deep learning one using U-Net model and the classical path using gradient image and watershed. Both those paths lead to segmented objects that allow us to find the center of the crystals.

For the detection branch, in this paper, we focus on two approaches: one relying on pre-processing the image using a partial differential equation that could somehow enhance the borders of the crystals and, therefore, approximate their center; and another one that uses machine learning (specifically the yolo model) to detect elements in the image.

Figure 1 shows those branches and their paths. In this work, we followed only the detection branch exploring its two approaches: partial differential equation pre-processing and object detection deep learning model. The segmentation path can provide well-established workflows, but the discussion is out of the scope of this work. Considering both chosen paths, the crystal identification for a single raw image is made of a sequence of algorithmic operations according to Figure 2, also indicated in the following table:

**Figure 1:**
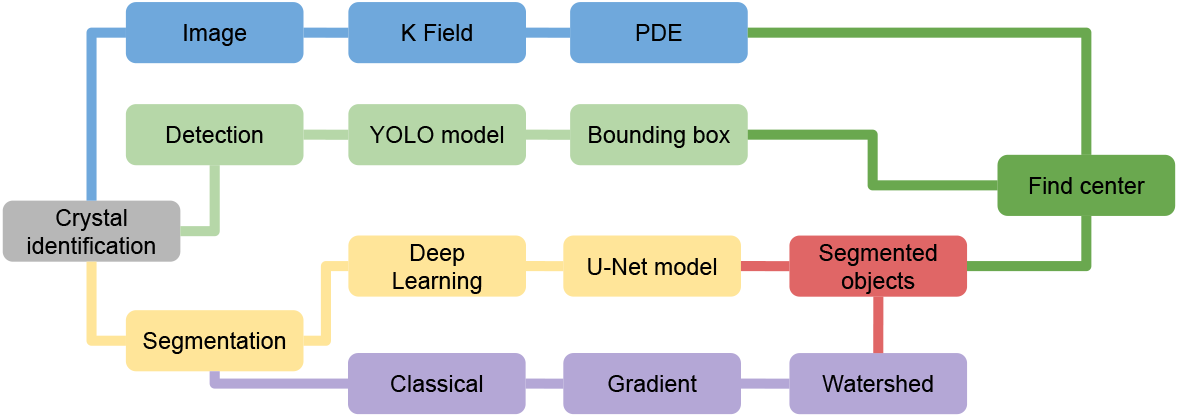
Proposed approaches presented on December 13th 2023. We chose to follow the detection branch and the segmentation branch is a proposed future work for this problem.

**Figure 2:**
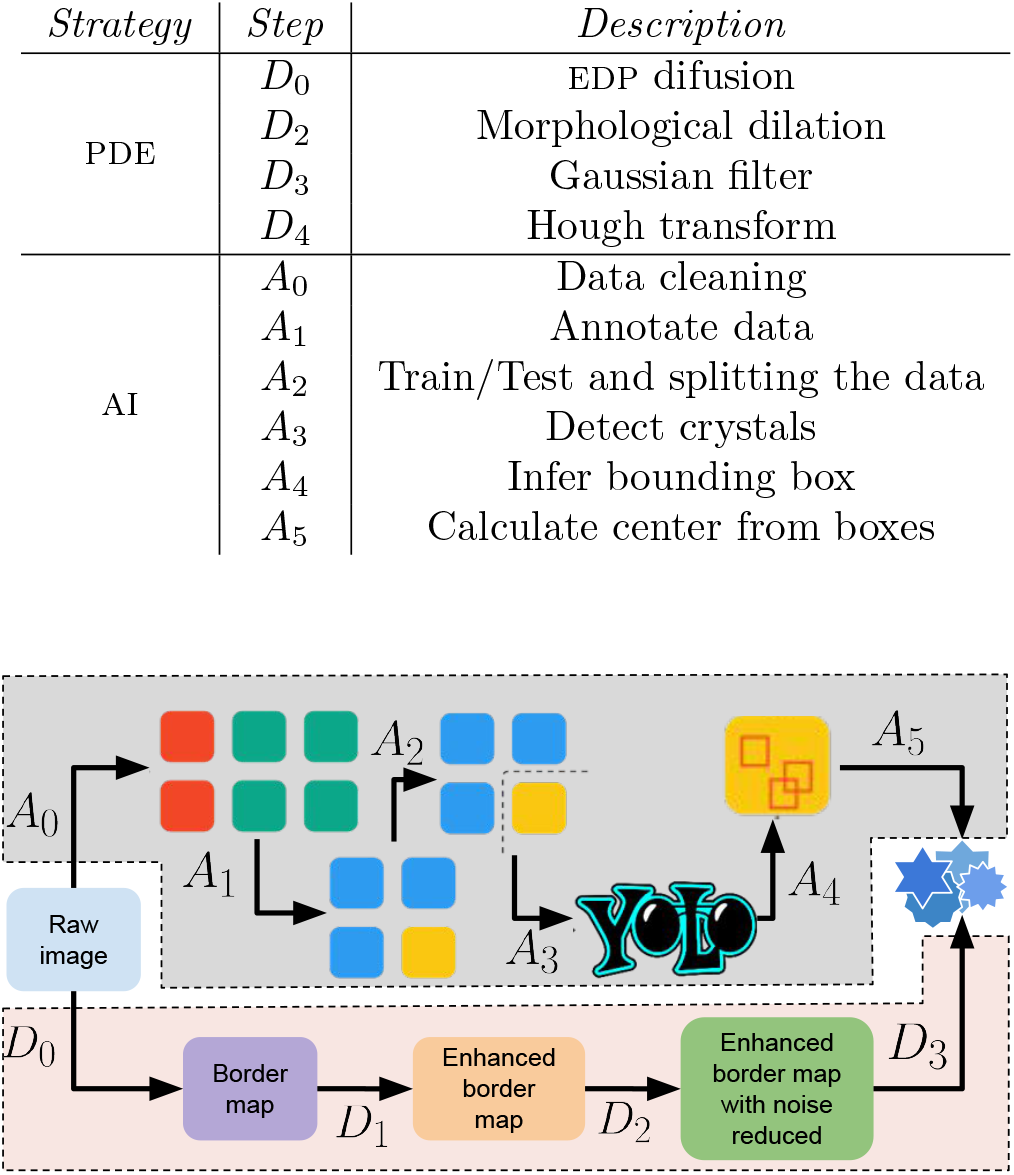
Two proposed and implemented strategies. The data cleaning and YOLO approach and the direct detection through the solution of a PDE and applying filters after that.

## 3 Evolution PDE

The partial differential equation model considered is given by a diffusion equation with homogeneous Neumann boundary conditions, as given by the following Eq.(3.1),

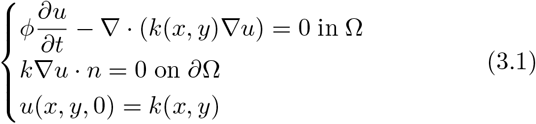

with *ϕ* = *k*(*x, y*). This model describes physical situations such as heat conduction, flow through a porous medium, and other processes that refer to transporting some substance. As mentioned above, we consider homogeneous Neumann boundary conditions, which means that the normal derivatives in the borders of the domain are zero, this states no flux in or out of the domain. The scalar field *k*(*x, y*) shown by (3.1) can be interpreted as a diffusion coefficient, determining how *u* flows along time in the domain.

The initial value for *u* is set as the scalar field *k*(*x, y*) itself. We can use an implicit Euler scheme for the temporal derivative and a central finite differences scheme for the spatial derivatives.

More details about the discretizations are in [6]. The main input for this model is the scalar field *k*(*x, y*), for the tests presented here, we used the grayscale raw image with values between 0 and 1. As for the parameters, users must set only the final time for the simulation, since we have an equation that varies over time. Given the implicit time discretization as in Eq.(3.2)

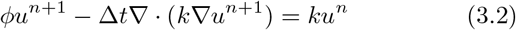

with Δ*t* being the time step, and *u*^*n*^ is the solution in time *t*_*n*_, we simulate until we reach *t* = *t*_*f*_ (*t*_*f*_ is the final time) and perform a discrete derivative in time, as in Eq.(3.3)

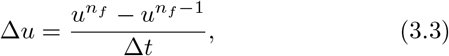

with *n*_*f*_ the final time step.

The scalar field Δ*u* is the one that we actually use as the border map. Since the image background almost does not vary over time, when we perform this discrete-time derivative, we somehow clean (at least a first cleaning) the background. Another parameter that can be explored is the *ϕ* function, for the tests presented here, we considered *ϕ* = *k*(*x, y*), but we can set *ϕ* as any other function of *k*(*x, y*). Choosing further functions *ϕ* is an interesting path to explore in future works.

With this border map given by the PDE model, a morphological dilation [7] was applied to enhance the borders. With this result, a Gaussian filter [7] can be applied to the resulting image to reduce noise. Some edges can be broken, also it is possible to have salt-and-pepper-like noises in the background, to deal with that, we apply a Median Blurring filter. After that, circular Hough transform [8] was applied giving the approximate center of the crystals. The parameters that can be obtained using the Hough transform are the minimum/maximum radius and border detection, and it is possible to give those parameters as input during the measurement by the user also, those parameters can vary substantially depending on the sample under investigation.

## 4 YOLO (You only look once)

In computer vision, the object detection task consists of, given an input image, determining bounding boxes containing a set of objects of interest. Machine learning (especially Deep Learning) approaches have strongly pushed the development of generic pipelines, that rely on annotating data by experts and fitting those models to those annotations. In the Deep Learning scenario, Girshick *et al*. [9] have adapted the use of ConvNets (Convolutional Neural Networks) to the context of object detection, by applying consecutive forward operations on specific regions of the image.

This process can easily become heavy to compute, especially in scenarios with multiple objects to detect. To improve the performance of object detection, [4] developed the yolo object detection architecture, which can detect several instances of objects in a single forward process, strongly increasing its throughput, thus showing as a promising alternative in time-constrained scenarios.

## 5 Results and Discussion

We start demonstrating the pde/direct methodology using some test images provided by the Manacá beamline [5]. The resulting image of this process (the scalar field Δ*u*) is a border map like the one shown in Figure 3.b); Both Figure 3.(c) and Figure 3.(d) show the morphological dilation and noise reduction by the Gaussian filter. Following the process, Figure 4.(a) shows the found centers on the raw image, we can see that at least 80% of the total number of crystals were detected. However, a quantitative analysis is still necessary to confirm those results. The action of Hough transform is to approximate the center of the crystals and it is presented in Fig.4.(b).

**Figure 3:**
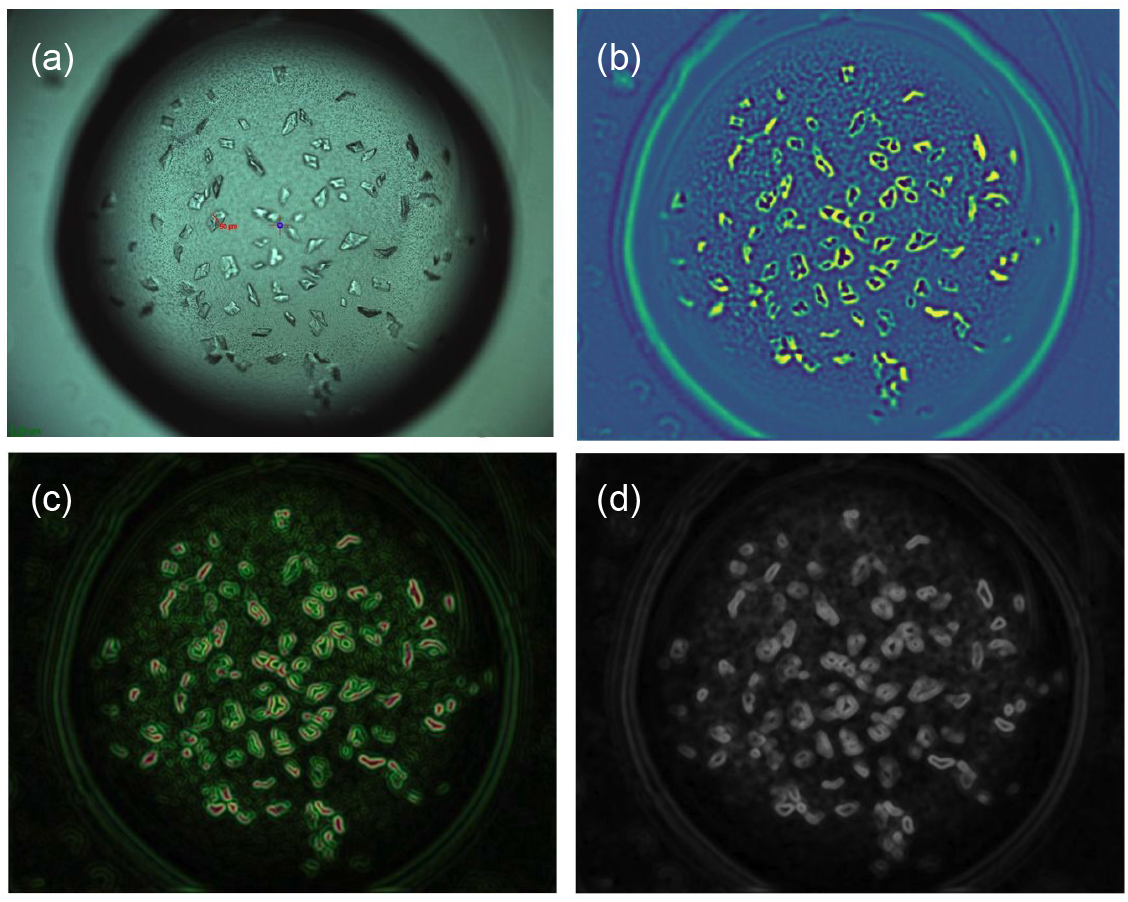
(a),(b) Raw image and border map after solving Eq.(3.1). The borders of the crystals are enhanced. (c),(d) Enhanced border map after the morphological dilation and noise-reduced image after applying the Gaussian filter.

**Figure 4:**
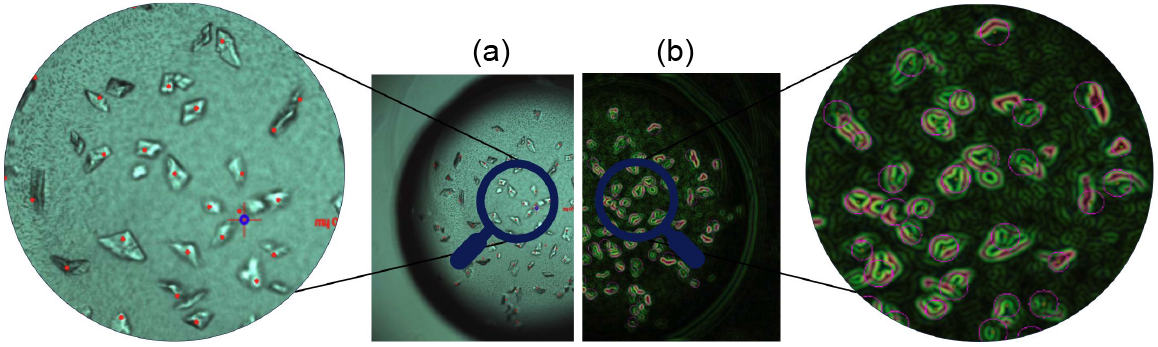
(a) Raw image with the detected centers. (b) Circles around the crystals after Hough transform.

For the Yolo approach, there are widely available models trained for detecting objects in natural images, however, they are not well suited to use in the crystallography scenario and thus require more fine-tuning. To properly adapt those models, we have manually annotated them using the online annotation tool MakeSense [10]. Due to the high similarity among samples, as multiple images are acquired from very similar parameters, we have manually selected a single representative from each scenario in a total of 43 available images, to be later presented to the model fitting.

A fine tune of 100 iterations was performed with the annotated samples in the YOLOv5l (previously trained in the COCO dataset) using an SGD optimizer with a fixed learning rate of 0.01. The best-performing model among all epochs was considered the final model. Even in the extremely constrained scenario, we achieved a reasonably well-fitted model with a great detection rate in a freshly acquired image never presented to the model (Figure 5).

**Figure 5:**
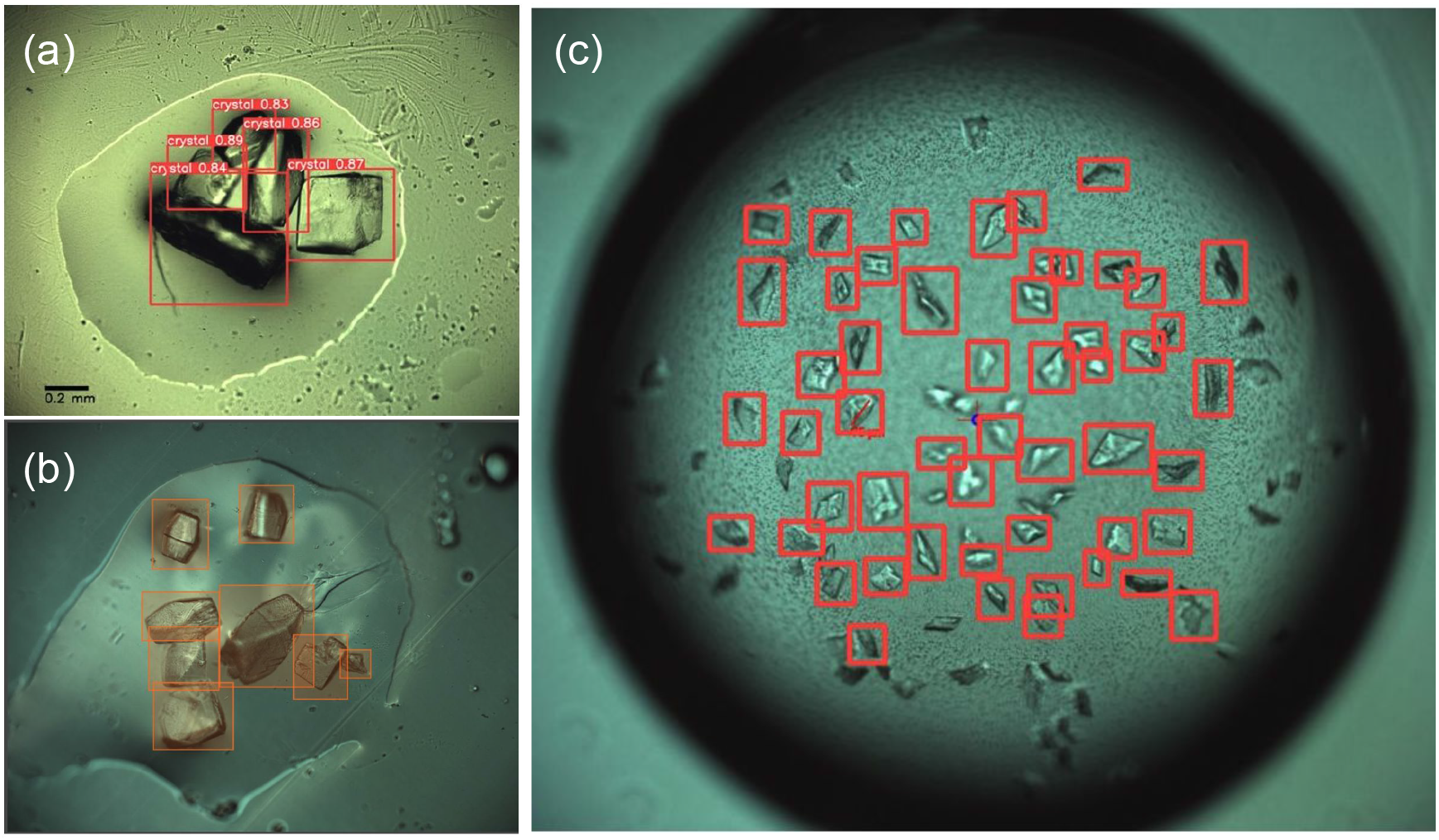
Our model applied in a dataset validation image (Left). Our model applied to an image acquired in a fresh experiment, never seen by the model.

## 6 Computational times

We computed the times for each approach (PDE and YOLO) and compared them in Table 1 shows the times for the PDE approach, *D*_0_ is the time for computing the Δ*u* field when solving the diffusion model. For the next steps: morphological dilation *D*_1_, gaussian filter *D*_2_ and Houg transform *D*_3_, since all of them are called by only one function in the pipeline and computed together giving the final result with the crystal’s centers, we computed the total time of the routine calling these steps.

**Table 1:** Computing elapsed times comparison between the pde approach and the yolo one. We used a standard image given by the beamline, with the usual resolution they will have in their operation.

| Strategy | Operation | Elapsed times (sec) |
| --- | --- | --- |
| PDE | $D_0$ | 6.4 |
| | $D_1, D_2, D_3$ | 0.6 |
| YOLO | $A_0, A_1, A_3$ | - |
| | $A_3, A_4, A_5$ | 2.5 |

Table 1 shows the times for YOLO approach, the steps *A*_0_ to *A*_2_ (data cleaning, annotating data, and testing/training the model) are pre-processing steps since they are related to training the model and we are not considering the time elapsed for this here since it can take up to some hours. We pre-trained a model given to the beamline so they can use it. Afterward, if necessary, they can train a new model. The remaining steps (*A*_3_ to *A*_5_ for detecting crystals, inferring bounding boxes, and calculating center from bounding boxes) are computed in one single routine in the pipeline, therefore we considered a single batch of time, giving the final result as the list of crystal’s centers.

## 7 Implementation

In the PDE approach, the finite difference discretization for the spatial derivative and the implicit Euler scheme for the time derivative generate a linear system *Ax* = *b* with *A* a pentadiagonal matrix [6]. For this implementation, we chose to use an iterative solver for the linear system, particularly the Gauss-Siedel algorithm, therefore, we did not need to compute and save the matrix *A*. This is a common choice when dealing with standard fluid mechanics simulations, especially when one knows that the problem behaves well enough and needs only a few iterations of the linear solver. That was the case in this work since the model is equivalent to a classical one for fluid mechanics, indeed, for the tests presented here, the Gauss-Siedel algorithm needed more or less 15 iterations to converge.

For each iteration of the system solver, we need to loop over all the image’s pixels. This is a time-consuming operation depending on the size of the image. To optimize this step, this loop was implemented in C. We did not use multi-threads or parallel programming, and the serial code in C was enough to accelerate the Python code for our purposes. Cython was used along with Python for this step.

## 8 Conclusion

We presented two approaches for finding crystals and their centers in images obtained by a microscope in the Manaca beam-line. This procedure is necessary for the beamline operation since it allows the scientists to measure the diffraction patterns of those crystals. Once they know the centers, they can move each crystal to the beam and realize the measure.

The approaches were: a diffusion PDE model to enhance the borders of the crystals followed by applying the Hough transform and using YOLO with a pre-trained model to detect crystals in the images. Comparing the times in Table 1, one can see that the YOLO approach took less time to find the centers if we do not count the time used to train the model, this procedure)could take hours depending on how many images are there to annotate and computational resources available.

The PDE approach took more computational time when compared to YOLO, but it is a direct method that does not require training or annotating. Also, it is possible to investigate methods to accelerate the approach: we can change the linear system solver, change the final time for the temporal loop, or even use parallel programming with tools like multithreads, MPI, or CUDA. There are also plenty of possibilities to investigate that can enhance the method for better detecting the crystals with some adjustments to the diffusion equation.

Overall, the computational times for both approaches were satisfactory for the beamline, which required at least one minute to compute the centers with 80% of crystals detected in the image. Both methods could detect at least 80% of the crystals for the images that we tested, more tests are now in course in the beamline using images that they are acquiring during operation.

## Acknowledgments

We would like to acknowledge the Brazilian Ministry of Science, Technology, and Innovation (mcti) for funding this work, through the Brazilian Center for Research in Energy and Materials (cnpem). We also thank to the 2023 Integrative Think Tank initiative, promoted by the University of Bath through Samba project (EPSRC Centre for Doctoral Training in Statistical Applied Mathematics) and held in the Sirius, the 4th generation Brazilian Synchrotron at CNPEM.

## Code Availability

We provide a computing package including all the codes described in this manuscript. The codes are mostly written in c and Python using other open resources such as OpenCV (smoothing images [11], morphological transformations [11], and features detection Hough circles [11]) and Cython. Since Python is one of the most used scientific programming languages throughout similar facilities, our package also includes Python bindings, allowing users with different architectures. The codes are made available under a gpl license hosted in Zenodo **(10.5281/zenodo.13361337)**. For distribution, the package name is ssc-fext (standing for feature extraction), where the acronym ssc stands for Sirius Scientific Computing, which is the computing efforts from lnls and the Data Acquisition and Processing Division for beamline support across sirius. An example for using the package is given by the Scripts 1 and 2.

**Script 1:**
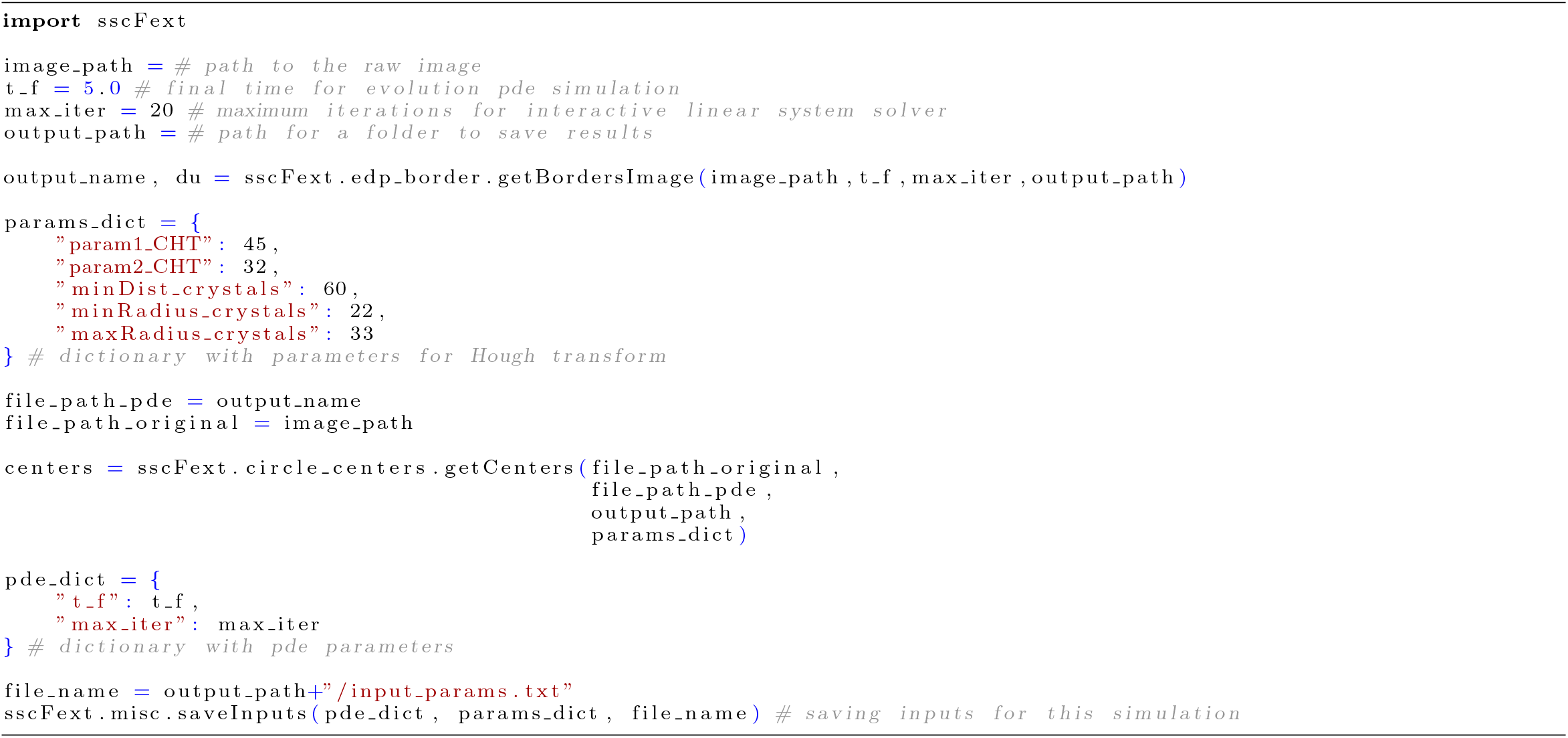
Example using package `ssc-fext` for PDE approach.

**Script 2:**
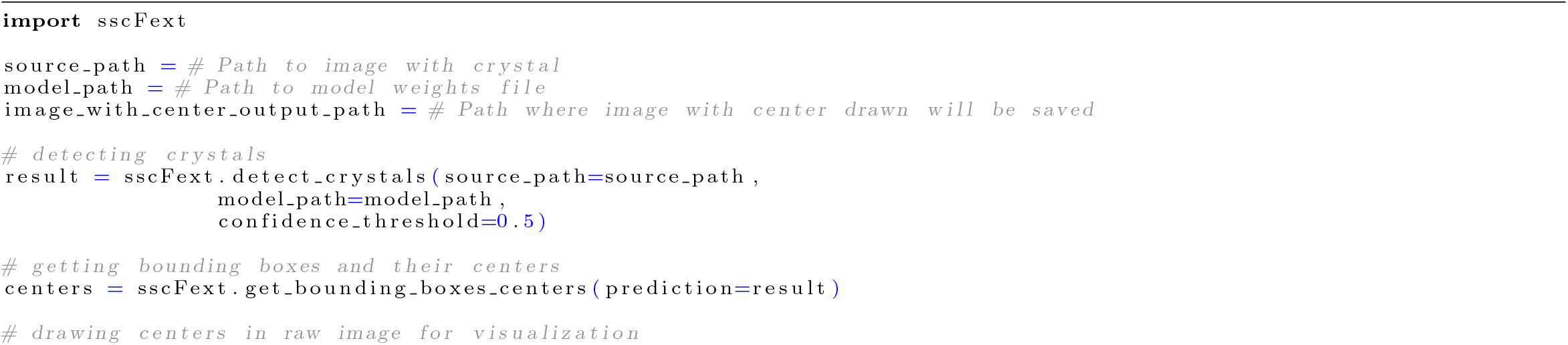
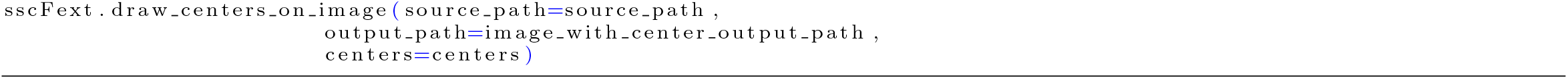
Example using package `ssc-fext` for YOLO approach, the model was trained before this process as explained earlier

## References

[1] Carmelo Giacovazzo. Fundamentals of crystallography, volume 7. Oxford university press, USA, 2002.

[2] Marcus Oscarsson, Antonia Beteva, David Flot, Elspeth Gordon, Matias Guijarro, Gordon Leonard, Sean McSweeney, Stephanie Monaco, Christoph Mueller-Dieckmann, Max Nanao, et al. Mxcube2: the dawn of mxcube collaboration. Journal of synchrotron radiation, 26(2):393–405, 2019.

[3] Itt (integrative think tank) 2023. https://lnls.cnpem.br/eventos/integrative-think-tank-brazil-2023/. Accessed: 2024-04-23.

[4] Joseph Redmon, Santosh Divvala, Ross Girshick, and Ali Farhadi. You only look once: Unified, real-time object detection. In Proceedings of the IEEE conference on computer vision and pattern recognition, pages 779–788, 2016.

[5] Andrey Nascimento, Evandro Araujo, Carlos Hagio, Silas Almeida, Ana Carolina Rodrigues, Daniela Barretto Barbosa Trivella, Marjorie Bruder, Joane Kathelen Rustiguel, Cristina Dislich Ropke, Lisandra Ravanelli Pessa, et al. Launch of the manacá beamline at sirius: First protein crystallography structures and new opportunities for pharmaceutical development using synchrotrons. Synchrotron Radiation News, 34(5):3–10, 2021.

[6] Randall J LeVeque. Finite difference methods for ordinary and partial differential equations: steady-state and time-dependent problems. SIAM, 2007.

[7] Rafael C Gonzales and Paul Wintz. Digital image processing. Addison-Wesley Longman Publishing Co., Inc., 1987.

[8] Richard O Duda and Peter E Hart. Use of the hough transformation to detect lines and curves in pictures. Communications of the ACM, 15(1):11–15, 1972.

[9] Ross Girshick, Jeff Donahue, Trevor Darrell, and Jitendra Malik. Rich feature hierarchies for accurate object detection and semantic segmentation. In Proceedings of the IEEE conference on computer vision and pattern recognition, pages 580–587, 2014.

[10] Make sense ai. https://www.makesense.ai/. Accessed: 2024-04-23.

[11] G. Bradski. The OpenCV Library. Dr. Dobb’s Journal of Software Tools, 2000.

